# FAIR enough? Status of metabarcoding sequence read data and metadata in public archives

**DOI:** 10.1101/2021.09.23.461561

**Authors:** Christiane Hassenrück, Julia Matela, Tobias Poprick, Véronique Helfer, Massimiliano Molari, Raïssa Meyer, Pier Luigi Buttigieg, David Riedinger, Peter Woollard, Frank Krüger, Ivaylo Kostadinov

## Abstract

Metabarcoding is well established across research fields for characterising biological communities. However, empirical evidence suggests that data (re-)usability remains limited. Here, we systematically assess the FAIRness of publicly available metabarcoding data, and the compliance of the sequence data files with submission requirements. While minimal mandatory metadata, such as geographic information and collection date, were retrievable for ∼90% of the sequencing runs, interoperability was not always ensured, and other essential metadata about the environmental origin and amplified marker gene were frequently neither provided nor syntax-compliant. Overall, with ∼30% of runs exhibiting inconsistencies in their metadata and more than half not linkable to a scientific article to retrieve further information, a large amount of existing data sets will likely remain non-reusable. Furthermore, less than 10% of the sequence data itself was submitted in the format required by the data archive. To tackle the most immediate needs to improve FAIR sequence data management, we provide a list of suggestions for researchers, research institutions, reviewers, publishers, brokers and databases, that we believe will have a large positive impact on sequence data FAIRness.

## Introduction

Next-generation sequencing is now firmly established as a routine tool in multiple fields of the life sciences, such as ecology (foremost microbial ecology), biodiversity research, and conservation biology. Knowledge derived from nucleotide sequence data, also referred to as digital sequence information (DSI), is becoming increasingly relevant for decision-making in natural resource management (e.g. Sustainable Development Goals^1^, Water Framework Directive^2^) and as part of international agreements (e.g. Convention on Biological Diversity^3^). The amount of nucleotide sequence data continues to grow exponentially^4^. However, a string of nucleotides (ACTG) alone does not contain much information - metadata and contextual parameters (Box 1) are required to describe sample origin and sequence generation for the data to be meaningful within and beyond the scope of the study, for which the sequence was obtained. Capturing and communicating not only the primary (sequence) data, but also its metadata and contextual data, is a crucial part of good data management. To promote sustainable data management and usage, the FAIR principles were introduced. They offer guidance on how to make data Findable, Accessible, Interoperable, and Reusable (Box 1)^5,6^, to prepare for a future of more automated analyses with the aim that the value of the data will not be restricted to a single study, but will extend to the reuse and integration across multiple studies over time. Growing volumes of data have resulted in nucleotide sequence data being used more frequently in conjunction with data science. This enables new scientific questions to be answered with existing data constituting a public good for the scientific community (Box 1). One prime example is the TARA Oceans data set^7^, which has already resulted in hundreds of publications, highlighting the value of secondary data use.

The implementation of metadata standards is a key aspect of FAIR data (Box 1)^6^. Already in 2011, the Genomics Standards Consortium (GSC)^8^ established the ‘Minimum Information about any (x) Sequence (<u>MIxS</u>)’ family of standards, to describe sample collection, processing, and sequence generation in a congruent manner (Figure 1)^9^. MIxS consists of several customised checklists and extensions tailored for a diverse set of sequencing applications and investigated environments, defining mandatory (core) and checklist-dependent recommended and optional parameters (metadata fields). In addition to standardising metadata field names, MIxS provides a consistent format for units and syntax for data values, thereby promoting the reusability of the data provided in compliance with this standard. Further, MIxS makes use of ontologies, such as the Environmental Ontology (<u>ENVO</u>)^10,11^ or the Experimental Factor Ontology (<u>EFO</u>)^12^, to describe the sampled environment or experimental conditions using a controlled vocabulary and to make the metadata more interoperable and machine-actionable. Since its adoption, several major new releases of MIxS have been published, updating metadata field names, syntax, requirement status (mandatory, recommended, optional), and extending the list of available checklists as new sequencing approaches become available.

**Figure 1:**
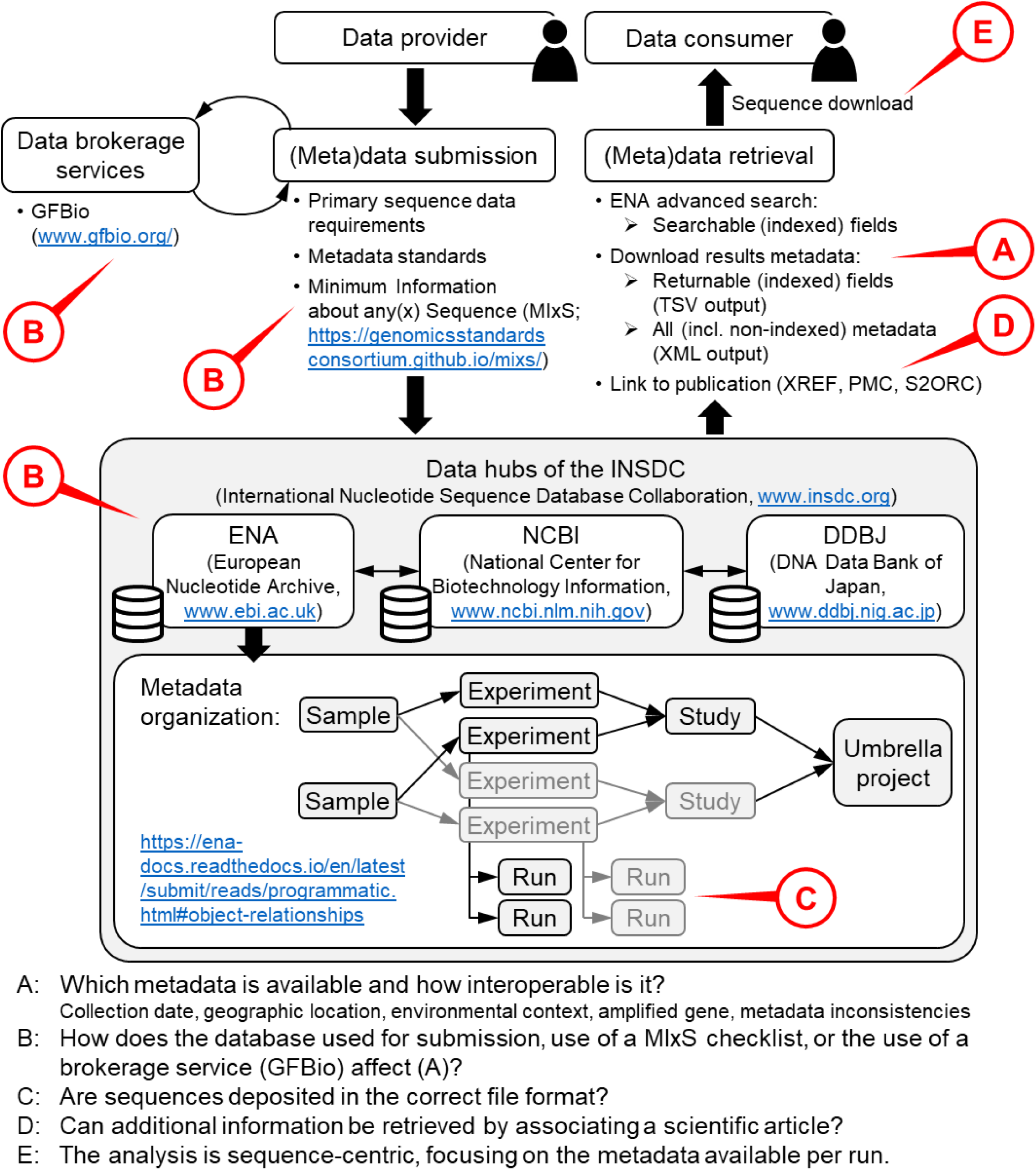
Summary of the submission and retrieval process for nucleotide sequence data using the databases of the INSDC. The letters highlight the specific aspects in that process that were investigated here, focusing on how metadata quality (A) is affected by various factors (B), the compliance of the primary sequence data with data archiving requirements (C), and the retrieval of further information via associated publications (D) in a sequence-centric data mining approach (E).

### Box 1: Glossary

**FAIR** (paraphrased after Wilkinson et al.^6^:

- Findable: Data and metadata are linked and findable via a unique and persistent identifier (e.g. accession number). Metadata is further searchable.
- Accessible: Data and metadata are retrievable (by humans and machines) via their identifier. Metadata remains accessible even if associated data is not available anymore.
- Interoperable: Data and metadata use a common language for knowledge representation understandable by humans and machines.
- Reusable: Data are described by rich metadata to provide the context required for reuse.

**Ontology:** Ontologies are formal representations of domain knowledge that define uniquely identified terms and logical relationships between them, enabling both human understanding and machine processing for data integration and interoperability across systems. In the case of the Environmental Ontology (ENVO), this increases the interoperability of environmental descriptions, helping (meta)data records achieve demonstrable FAIRness.

**Metadata:** Data that describe the primary data. Here, primary data are nucleotide sequence data. Most metadata parameters are intrinsic to the sampling or experimental design and the laboratory or analytical procedures. As such they are often known *a priori*, i.e. before the primary data collection. For instance, sampling location, experimental treatments, sequencing platform.

**Contextual data:** While often grouped together with metadata, contextual data is referring to parameters which are recorded alongside the primary data. For instance, temperature, salinity, inorganic nutrient concentrations. They are often primary data for other research fields.

**Metadata standards:** Metadata standards provide a structured framework for metadata documentation in compliance with the FAIR principles. They provide checklists of well-defined parameters, also referred to as metadata fields, to be reported and determine the vocabulary and syntax to be used to ensure consistent data across studies.

**Data mining:** In the context of this publication, data mining is referring to the retrieval and reuse of data sets that have been archived in open access data repositories.

The main hubs for long-term nucleotide sequence data storage are the resources making up the International Nucleotide Sequence Database Collaboration^13^ (INSDC; Figure 1): the European Nucleotide Archive (ENA) at the European Molecular Biology Lab (EMBL) - European Bioinformatics Institute (EBI), GenBank and the Sequence Read Archve (SRA) at the National Center for Biotechnology Information (NCBI), and the DNA Databank of Japan (DDBJ) at the Research Organisation of Information and Systems (ROIS) - National Institute of Genetics (NIG). These databases mutually mirror to enable the same data coverage among them. The establishment of this infrastructure has been instrumental in propagating global standards for sequence data and metadata (e.g. fasta, and fastq data formats) and it offers services far beyond the provision of access to such data^14–16^. Furthermore, as part of the GSC, the members of the INSDC have contributed to designing the MIxS standards, pioneering the implementation of such metadata standards in nucleotide sequence databases with their official adoption in 2011^9^. For instance, while minimal specifications for sequence submissions, consisting of four mandatory metadata fields (collection date, country, a unique sample name/alias, and the NCBI taxon ID), have been agreed upon across the INSDC, each member recommends extensive <u>sample</u> <u>metadata checklists</u>^17^ substantially based on MIxS, which contribute to metadata field and value alignment. However, despite their basis in MIxS and regular updates to accommodate newer versions of MIxS, these checklists do not represent a 1:1 implementation, differing in syntax rules and requirement status.

To accurately represent metadata for next generation sequencing data along the process from sample collection to data generation, it is organised on several levels (Figure 1): **Runs** hold the sequence data generated by the sequencer. They are associated with **experiments**, which hold the metadata for individual library preparations and how these libraries were sequenced. Depending on the experimental set-up, they may also represent different nucleic acid extractions from the same sample. Experiments are collected into **studies**, which may use a common methodological approach, such as metabarcoding of a particular marker gene. Several studies can be summarised by an umbrella project that may correspond to the larger scientific project, for which the data was generated. **Samples**, referring to the biological material, most often its nucleic acid extract, can be associated with multiple studies through different experiments. This flexible metadata model allows representing complex experimental set-ups correctly, but can be hard for inexperienced data submitters to navigate properly and provide the necessary and standards-compliant information. This process is further complicated by the fact that MIxS checklists are implemented at the sample level, although they contain metadata fields more appropriate at the experiment level, such as ‘target_gene’, which may take on different values if sequencing libraries were prepared from the same nucleic acid extract.

When accessing sequence data as data consumer (Figure 1), all provided metadata for each level (run, experiment, study, sample) can be retrieved in XML format. To simplify metadata access, the <u>ENA advanced search</u>, aka the ENA Portal API^18^) offers a collection of indexed metadata fields that use standardised names and are searchable (i.e. usable to restrict the search) and returnable (i.e. downloadable in a user-friendly TSV format; Figure 1). At the run level, these indexed fields are also inherited from sample and experiment metadata. Currently, the implementation of indexed fields is limited to metadata that are mandatory for most checklists and/or most frequently provided. As such, many recommended and optional MIxS parameters are not indexed and only accessible in XML format, where no standardised nomenclature, controlled vocabulary or syntax are enforced. Therefore, some of the value of MIxS is essentially lost, making non-indexed metadata fields not interoperable.

In addition to metadata requirements, ENA also standardises the format of the submitted sequence data depending on the sequencing approach. For instance, paired-end Illumina raw reads have to be submitted as demultiplexed R1 and R2 files (fastq) without artificial sequences (e.g. adapters, linkers, barcodes/tags, primers) and prior to any quality trimming^19^. To provide sequencing data in such a format, initial sequence processing steps are necessary, starting from the multiplexed sequencer output. As bioinformatic sequence analysis pipelines vary, it is important to not deviate from the sequence format requirements by adjusting analysis workflows accordingly.

To support sustainable data management, data brokerage services, such as the German Federation for Biological Data (<u>GFBio</u>), have been established^20^. Brokerage services offer a central entry point for data submissions, providing personal guidance (helpdesk) on FAIR data, supporting and often simplifying the data submission process, and ensuring data deposition on the most appropriate archive.

To facilitate data reuse in data mining endeavours, access to and retrieval of the raw read sequencing data and metadata at the run level, together with the inherited metadata fields describing the sample and sequencing experiment, are most crucial. However, despite the available framework for FAIR data archiving, data interoperability and reusability are still limited and often complicated by insufficient metadata^21–23^ (personal observation). Therefore, we decided to conduct a review of the status of nucleotide sequence data and associated metadata accessible through ENA to (i) identify deficits in metadata quality and (ii) provide suggestions for improving FAIR sequence data management. Specifically, we investigated the following aspects (Figure 1):

- Interoperability of the indexed metadata (i.e. those returnable in TSV format by the ENA portal API), focusing on their availability, adherence to syntax requirements according to submission checklists, and the correct use of ontologies and controlled vocabulary
- Accessibility to and availability of additional information retrievable in the form of natural text, such as the non-indexed metadata fields provided in the XML format and scientific articles, in which the data is described
- The effect of MIxS usage, submission through a broker, and the database, through which the data was submitted, on metadata quality
- Feasibility and reliability of the identification of a sample as environmental based on its metadata
- Compliance of the primary sequence data with formatting requirements

## Methods

For the assessment of metadata quality on the Sequence Read Archive (SRA) we selected a widely used approach from biodiversity research: paired-end metabarcoding, i.e. amplicon sequencing, of environmental samples on the ILLUMINA platform. To identify such data sets, we constructed the following search query via the ENA Advanced Search to retrieve metadata at the level of raw reads (date accessed 2025-09-01):

tax_tree(12908) AND (instrument_platform=“illumina” AND library_layout=“paired” AND library_strategy=“AMPLICON” AND library_selection=“PCR” AND ( library_source=“genomic” OR library_source=“metagenomic” ))

With the NCBI taxon ID 12908, we aimed to retrieve all unclassified sequence entries, including their lower taxonomic ranks, such as ecological and organismal metagenomes. The combination of library strategy (“AMPLICON”), library selection (“PCR”), and library source (“METAGENOMIC”) should be indicative of metabarcoding data, i.e. reads generated from a specific marker gene amplified via PCR from a mixed, i.e. metagenomic, DNA template. The search query was further extended to also include “GENOMIC” as library source, since it was frequently used for metabarcoding studies in the scientific literature. This search returned metadata for 4,844,334 runs, which were downloaded in TSV format. To avoid biasing our results due to differing data practices across different fields, any search results indicating human origin and from clinical settings were removed, as different best practices for data sharing related to data protection and privacy may apply. This filter was implemented using the NCBI taxon ID, i.e. the scientific name, removing those which contained the keywords “human” and “enrichment culture” or were part of a manually curated blacklist (https://github.com/chassenr/ENA-metadata). For the remaining 3,461,316 runs, 3,455,898 experiments, 2,986,834 samples, and 44,421 studies, we further downloaded all available metadata at experiment and sample level in XML format to extract additional metadata fields attributes that were not parsed in the TSV output.

To identify the scientific articles associated with the sequencing data as potential source of additional metadata, we used the ENA XREF API with the primary and secondary study accessions. Furthermore, the PubMedCentral^24^ (version of 2025-09-02) and Semantic Scholar Open Research Corpus^25^ (version of 2025-08-19) were downloaded and searched for the regular expression corresponding to the primary and secondary study accessions^26^ of the sequencing data.

The availability and interoperability of the following metadata fields was evaluated relying mostly on the implementation of the MIxS checklists on ENA:

- Collection date: Values are expected to comply with the international standard ISO 8601 (*_date) or use a controlled vocabulary for missing values^27^ (“*_missing”). All remaining values were labelled as “*_other”. If no values were included in the TSV output (“tsv_*”), suitable metadata fields in the sample XML were screened (“xml_*”). Runs without collection date values were labelled with “no”.
- Geographic coordinates: Latitude and longitude values should be formatted as decimal degrees with a maximum of 8 decimal digits (“*_dd”) or use a controlled vocabulary for missing values (“*_missing”). All remaining values were labelled as “*_other”. If no values were included in the TSV output (“tsv_*”), suitable metadata fields in the sample XML were screened (“xml_*”). Runs without any latitude or longitude values were labelled with “no”.
- Country: Values should come from the <u>INSDC country list</u> (“*_country”) or use a controlled vocabulary for missing values (“*_missing”). All remaining values were labelled as “*_other”. If no values appear in the TSV output (“tsv_*”), suitable metadata fields in the sample XML were screened (“xml_*”). Runs without country values were labelled with “no”.
- Vertical coordinate: Values for depth, elevation, or altitude should comprise of a number but may contain additional elements (“*_number”). While we did not systematically assess this, values should be given in meters, with the unit not included as part of the value. Alternatively, the controlled vocabulary for missing values may be used (“*_missing”). All remaining values were labelled as “*_other”. If no values were included in the TSV output (“tsv_*”), suitable metadata fields in the sample XML were screened (“xml_*”). Runs without any depth, elevation or altitude values were labelled with “no”.
- Environmental context: The metadata fields environmental_sample (binary classification), tag, and the MIxS fields broad_scale_environmental_context, local_environmental_context, and environmental_medium should provide information about the environmental origin of a sample. The values for tag are automatically derived from the NCBI taxon ID (env_tax) and geographic coordinates (env_geo) to indicate the type of environment a sample originated from. Here, we adapted the confidence scheme of O’Cathail et al.^28^ to assign the labels “high”, “medium”, and “low” confidence, and “conflict” to each tag value. Values for the MIxS fields should follow the Environmental Ontology (ENVO) using the syntax “ocean biome [ENVO:00000015]” (“syntax”). If the value was either an element of an ENVO id or name (e.g. “ENVO:00000015” or “ocean biome”), it was labelled “name_id”. If the value could be approximated to an ENVO name after conversion to lower case alphanumeric characters, it was labelled “approx”. No similarity to ENVO names received the label “filled”. Runs without values were labelled with “no”. As such, we only assessed syntax and not semantic suitability of the provided metadata values.
- Amplified gene: The TSV output contains the column target_gene, which is extracted from the sample-level metadata, and the column library_construction_protocol parsed from the experiment-level metadata (“tsv_*”). Theoretically, information about the amplified gene is a feature of the library preparation and should therefore be expected at experiment level. We therefore screened suitable metadata fields of the experiment as well as the sample XML (“xml_*”) if no values were included in the TSV output. We distinguished between values that contained nucleotide sequences (“*_primers”) and those without (“*_other”). Runs without values were labelled “no”.
- Contradictory metadata: We investigated whether metadata fields contained values which are mutually exclusive in their meaning in the following three cases: (1) If a NCBI taxon id (scientific name) for a metagenome is provided, the library source should be given as “METAGENOMIC” and not as “GENOMIC”. (2) Paired-end sequencing data should be available in exactly two fastq files. (3) Although we restricted our search query based on the library selection, at the time of the search the ENA portal API was not implementing its own controlled vocabulary, but allowed partial string matches, so that “RANDOM PCR” and “RT-PCR” were also returned, which we used to identify inconsistencies. If sequences were generated from (meta)genomic material with the library strategy “AMPLICON”, it follows that the library selection must be ”PCR”. If either of these three conditions was not met, runs were labelled as “inconsistent (meta)data”, otherwise as “no inconsistencies found”.
- Discoverability of associated scientific article: Using a sequential approach based on the ease with which a scientific article could be associated with a sequencing data set, we labelled data sets as “xref_found” (ENA XREF API), “pmc_found” (regular expression search of the full PMC body of articles), or “s2orc_found” (regular expression search of the full S2ORC body of articles). Data sets for which no publication could be found were labelled as “no”.

Finally, overall metadata quality was determined based on the availability and interoperability of all the metadata fields assessed above:

- Machine-readable: Syntax-compliant metadata accessible in TSV format without inconsistencies. Metadata in this form will facilitate automatic data reuse.
- Human-readable: Metadata available in either TSV or XML format. Although these records are not syntax-compliant and may contain inconsistencies, they are linked to an associated scientific article for further information extraction. While data reuse may require manual curation, there is likely sufficient metadata available.
- Missing/inconsistent: Metadata is either missing or inconsistent and no scientific article can be found to validate and complete the metadata. This means that information crucial for data reuse is not available, potentially resulting in the loss of the data for future research.

We further evaluated the impact of the following factors on metadata quality:

- MIxS usage: We identified whether the MIxS standard was used for data submission based on mentions of MIxS and the selection of an ENA MIxS checklist according to the metadata fields “ncbi_reporting_standard” and “checklist”, respectively.
- Entry DB: The database, through which the sequencing data were submitted, was identified based on the syntax of the run accession^26^.
- Brokerage service: To evaluate the impact of using a (meta)data brokerage service, we scanned for the use of GFBio (the most frequently used broker across data sets) in the metadata field “broker_name”.

Their effect was quantified using chi-square contingency tests, followed by post-hoc tests implemented in the R package chisq.posthoc.test^29^ if applicable. Bonferroni adjustment was applied to all p-values to account for the multiple tests conducted for each factor.

Apart from metadata quality, the reusability of nucleotide sequence data - especially the automation thereof - strongly depends on the availability and format of the submitted read data itself. Therefore, we checked if a run accession actually contained data based on non-zero values in the metadata fields “base_count” and “file_size”. Furthermore, we used experiences from two data mining efforts focused on 16S rRNA gene metabarcoding data^30,31^ as case studies to evaluate if the raw read data (fastq files) were compliant with SRA requirement as outlined in the ENA submission documentation^19^. Specifically, we reported on the state of pre-processing (demultiplexing, adapter clipping, primer clipping, paired-end read merging) that the reads were in, considering that paired-end Illumina reads should be archived as demultiplexed, unmerged forward and reverse reads, without artificial sequences and prior to any quality trimming.

The analysis was implemented R version 4.4.0^32^ and python version 3.12.2^33^ supplemented by custom bash scripts, GNU parallel 20210822^34^, and jq version 1.6^35^. All scripts and code for the metadata assessment are available on: https://github.com/chassenr/ENA-metadata.

## Results

### General trends

The number of sequencing runs, for which metadata is accessible via the above-mentioned search query, has been increasing continuously over the last decade, with 30% of the runs and 43.5% of the data volume submitted since 2024-01-01 (Figure 2). The majority of runs were contributed through NCBI (85%), followed by ENA (12%) and DDBJ (2%). In total, 39% of runs have been submitted using a MIxS-related checklist or NCBI reporting standard. This percentage has been largely stable since 2020. Interestingly, a significantly larger fraction of runs submitted through ENA and DDBJ used a MIxS checklist compared to NCBI data submissions. The fraction of runs brokered by GFBio was comparatively low (0.5%) and fluctuated over the years without a clear increasing or decreasing trend ever since the launch of the brokerage service in 2016.

**Figure 2:**
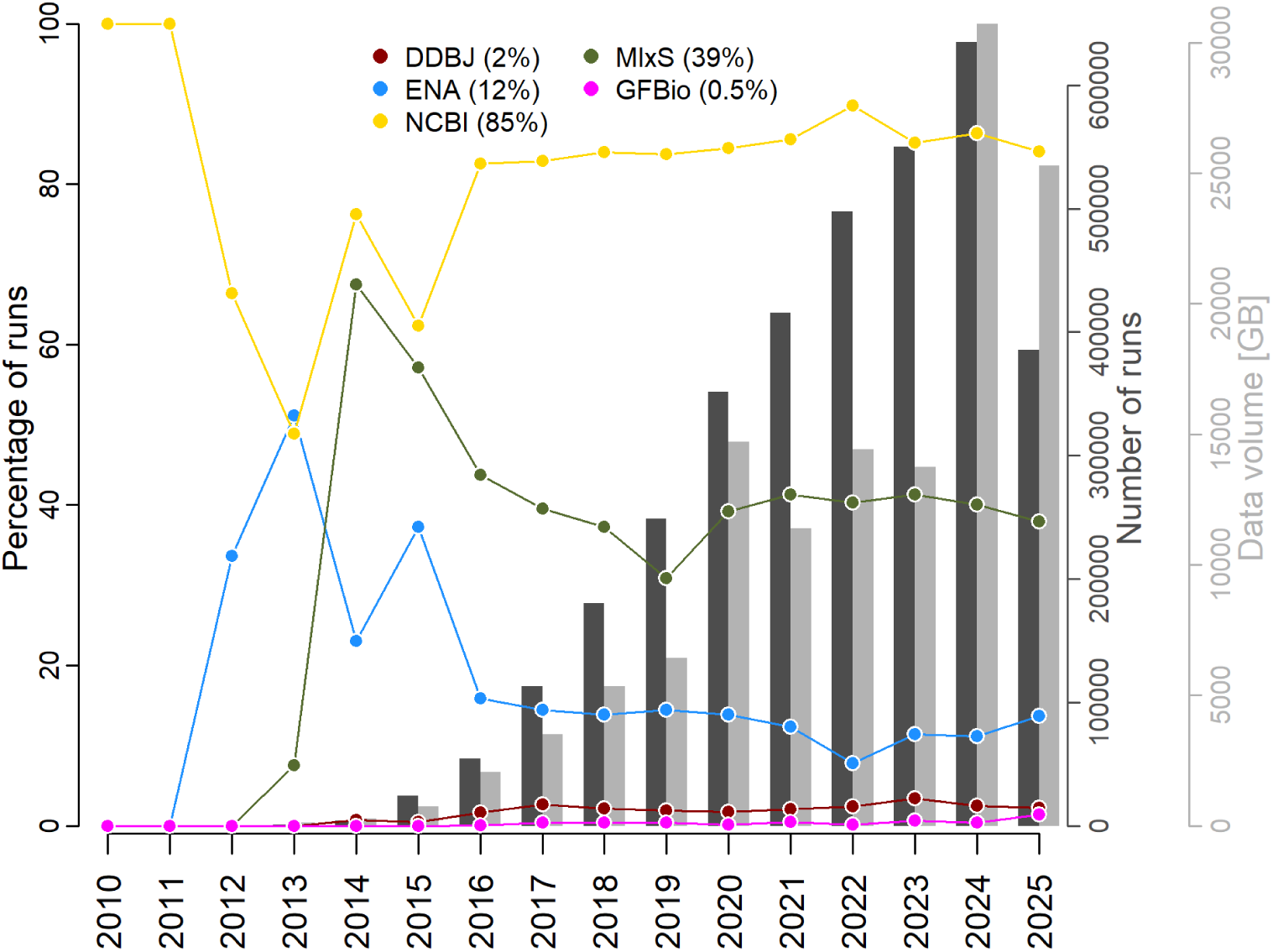
Overall trends in metabarcoding data submissions. Bars indicate number of runs and their data volume submitted per year (right axis), while lines follow the percentage of runs (left axis) submitted through each of the three member databases of the INSDC, those submitted using a MIxS checklist, and those whose submission was brokered by GFBio. Percentages in parentheses were calculated over all years. Note that for 2025 only data sets until 2025-09-01 are included.

On average 1.16 runs were submitted per sample with a highly skewed distribution, where only 7% of the samples were associated with more than one run. According to the definition of metadata object relationships, this would mean that for 93% of the samples, only 1 marker gene was sequenced. However, manual checks showed that there are studies, where samples processed for more than one marker gene were assigned independent accessions (PRJEB55296^36^), or physically independent samples were assigned the same sample accession number (SAMEA117556368^37^; SAMN23377276^38^).

Among the 25 most frequent NCBI taxon IDs (Supplementary Figure 1), were those of soil, plant, marine, freshwater, rhizosphere, root, seawater, wastewater and sediment metagenomes, suggesting that our filter was successful in largely capturing environmental metabarcoding data. However, other taxon IDs, such as metagenome (not further specified) and mouse gut metagenome, were more ambiguous. Indeed, they were not used to derive the values for “env_tax” in the metadata field “tag”. Despite this, since an environmental origin cannot be excluded, those data sets were kept for the further analysis.

### Collection date and geographic location

Here, we investigated metadata describing location in four dimensions (xyzt): longitude, latitude, vertical, and time. This metadata is particularly crucial to place an environmental data set in time and space and to derive further information from such coordinates, e.g. from gridded data (remote sensing data products, models).

Syntax-compliant values for “collection_date” (including the metadata fields “collection_date_start” and “collection_date_end”) were provided in the correct metadata field (i.e. returnable in TSV format) for more than 90% of runs, with an additional 4% using accepted syntax for missing values (Figure 3A). Since 2023, when “collection_date” became a mandatory metadata field for all data submissions even using minimum checklists (e.g. <u>ERC000011</u>), this percentage has even exceeded 98%. Only for a negligible fraction of runs (0.08%), for which the field “collection_date” was empty, some kind of ISO8601 compliant date was found in the sample XML. The use of a MIxS checklist or GFBio as brokerage service, significantly increased the number of runs with syntax-compliant collection date information, while reducing the number of runs without any date information. Interestingly, data sets submitted through NCBI also exhibited higher syntax-compliance (Supplementary Table 1). Despite the majority of collection date information being syntax-compliant and easily accessible in TSV format, not all such metadata may be actionable. For more than 30% of the runs, only year or month information was available under “collection_date” without a concrete day or range being provided. While, in theory, it is possible that samples spanning a longer collection period were pooled for sequencing, we rather suspect that independent samples were not declared as such. This suspicion was supported by more samples without concrete day information having a larger number of runs derived from them compared to those with concrete day information. Furthermore, the use of GFBio as brokerage service, i.e. a human checking the submitted metadata, drastically reduced the proportion of runs without concrete day information.

**Figure 3:**
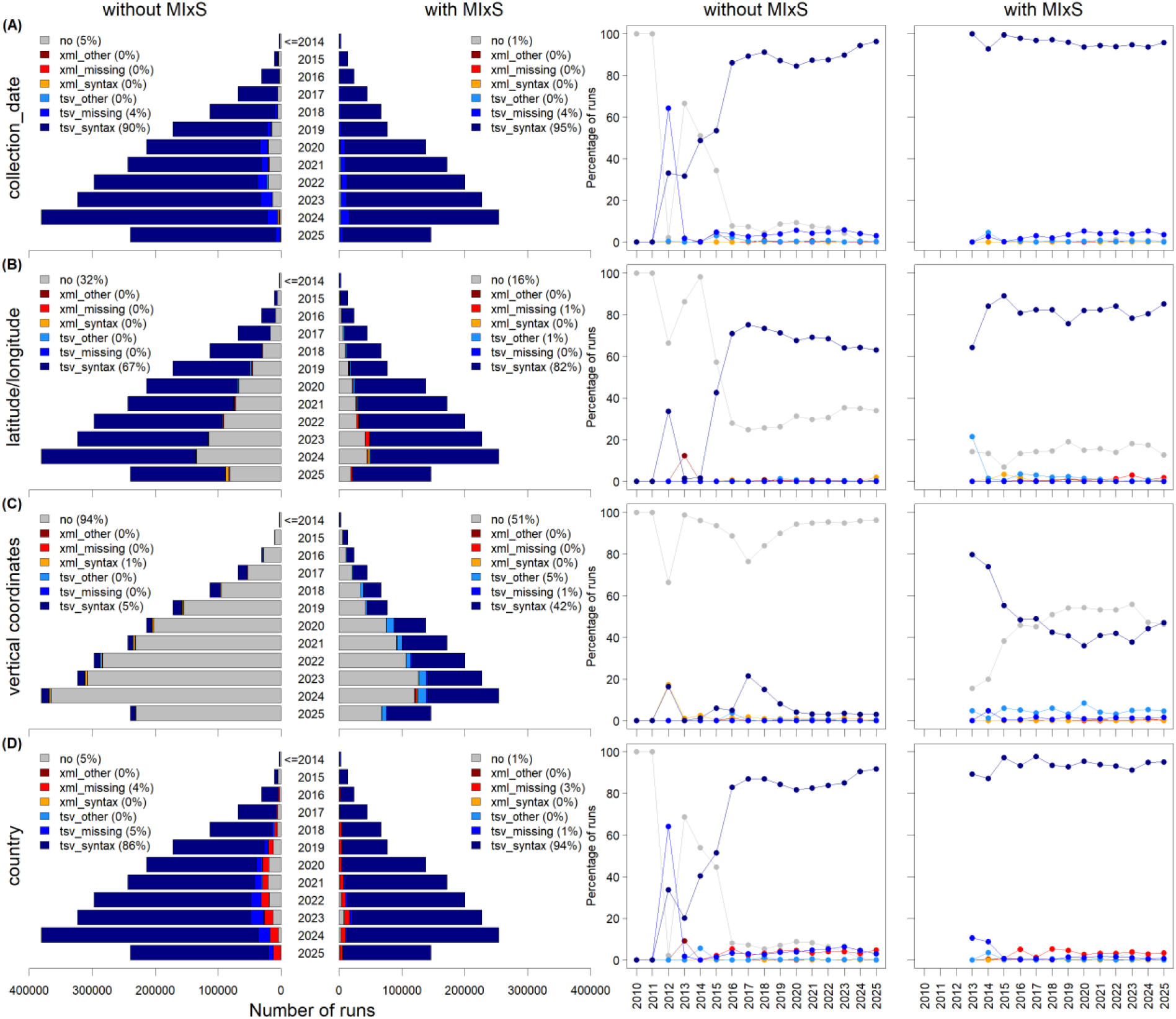
Availability and syntax compliance of (A) collection date, (B) latitude and longitude, (C) vertical coordinates (depth, elevation, altitude), and (D) country. The bar charts show the number of runs per submission year, separately for runs submitted without and with a MIxS checklist and coloured by metadata quality category. The line charts show the percentage of runs in each category over time. Categories are suffixed according to syntax compliance: *_syntax (syntax-compliant), *_missing (controlled term for missing values), *_other (free text, non-compliant value); and prefixed according to metadata accessibility: tsv_* (value available in the TSV search output), xml_* (value only available in metadata XML); with “no” indicating that no value was provided. Percentages in the legends refer to all years combined. Runs submitted before 2015 are pooled (≤2014) in the bar charts.

Latitude and longitude are mandatory environmental MIxS checklist fields. However, these metadata fields are not necessarily enforced outside the use of a MIxS checklist. Nevertheless, in our example the majority of all runs (73%) were archived with latitude and longitude available syntax-compliant as decimal degrees in the TSV search output (Figure 3B). No entries were found for syntax-compliant missing data. Searching the sample XML for putative latitude and longitude values also did not yield many results with only an additional 0.35% of all runs providing decimal degrees as part of non-indexed metadata fields. The use of a MIxS checklist reduced the percentage of runs without any latitude and longitude coordinates by half and increased syntax compliance for both indexed and non-indexed metadata fields, while the effect of using GFBio as brokerage service was more targeted through a significant increase in the provision of syntax-compliant decimal degree values returnable in the easily accessible TSV format (Supplementary Table 1). While data sets submitted through NCBI tended to more often contain syntax-compliant decimal degree values in the TSV output, richer metadata was found in the sample XML of data sets submitted through ENA in cases where no TSV-parsable values were provided.

To pinpoint a sample in space, information about its vertical position is often equally as important as its latitude and longitude coordinates. Altitude and elevation both express the distance relative to a fixed reference (e.g. sea level) either in the air or on a surface, respectively, while depth is measured relative to its closest surface. Depending on the MIxS environment-specific extension, these metadata fields are listed as recommended or mandatory. Our results showed that submissions without the use of a MIxS checklist hardly ever provided information about vertical coordinates, whereas 42% of the runs submitted using a MIxS checklist included syntax-compliant and TSV-parsable numeric values for at least one of the three metadata fields (Figure 3C). However, syntax requirements for altitude, elevation, and depth are very permissive (they only have to contain a number) and for 52% of the runs with syntax-compliant numbers additional characters were part of the value. This was not the case for data sets brokered by GFBio. There, syntax-compliant numeric values returnable in TSV format were available for 98% of the runs and only consisted of a number and/or range without non-numeric characters. Overall, if a syntax-compliant numeric value was provided together with a text string, those were corresponding to commonly used distance units (“cm”, “m”, “mm”, “inches”, “in”, “meters”, “meter”, “cmbf”, “mbsl”) in 92% of the runs. Regarding the effect of the entry database, submissions through DDBJ and ENA tended to provide information on vertical coordinates more frequently compared to NCBI (Supplementary Table 1).

Unlike latitude and longitude, the metadata field “country” is mandatory for all sequence read submissions, together with “collection_date”. As such, “collection_date” and “country” show comparable patterns in terms of availability, syntax compliance, and ease of access (Figure 3D, Supplementary Table 1).

### Description of sampled environment

The metadata fields “broad_scale_environmental_context”, “local_environmental_context”, and “environmental_medium” use ENVO terms to describe various characteristics of the sample and the environment it originated from. They are mandatory parameters for any MIxS checklist. As such, it is not unexpected that more than 90% of runs submitted according to a MIxS checklist provide a value for these metadata fields, while 96% of the runs without MIxS did not (Figure 4A-C). However, even then, ENVO terminology was inconsistently used with only 9 and 20% of runs meeting the MIxS syntax requirements for “broad_scale_environmental_context” and “local_environmental_context”, respectively. Runs submitted through ENA showed a higher frequency of syntax-compliant values compared to those submitted through DDBJ and NCBI (Supplementary Table 1). Furthermore, it is promising that the fraction of syntax-compliant values for those two metadata fields has been increasing since MIxS checklists were first used in 2013, from 0 – 11% (broad_scale_environmental_context) and 0 – 28% (local_environmental_context). Interestingly, for the metadata field “environmental_medium” the majority of runs (66%) displayed syntax-compliant values if provided. Furthermore, the fraction of runs with MIxS checklist usage, for which the values for “broad_scale_environmental_context”, “local_environmental_context”, and “environmental_medium” could be coerced to resemble ENVO term names and/or IDs by conversion to alphanumeric, was above 80%, suggesting rudimentary interoperability. Runs submitted through GFBio were almost always syntax-compliant for those metadata fields (Supplementary Table 1).

**Figure 4:**
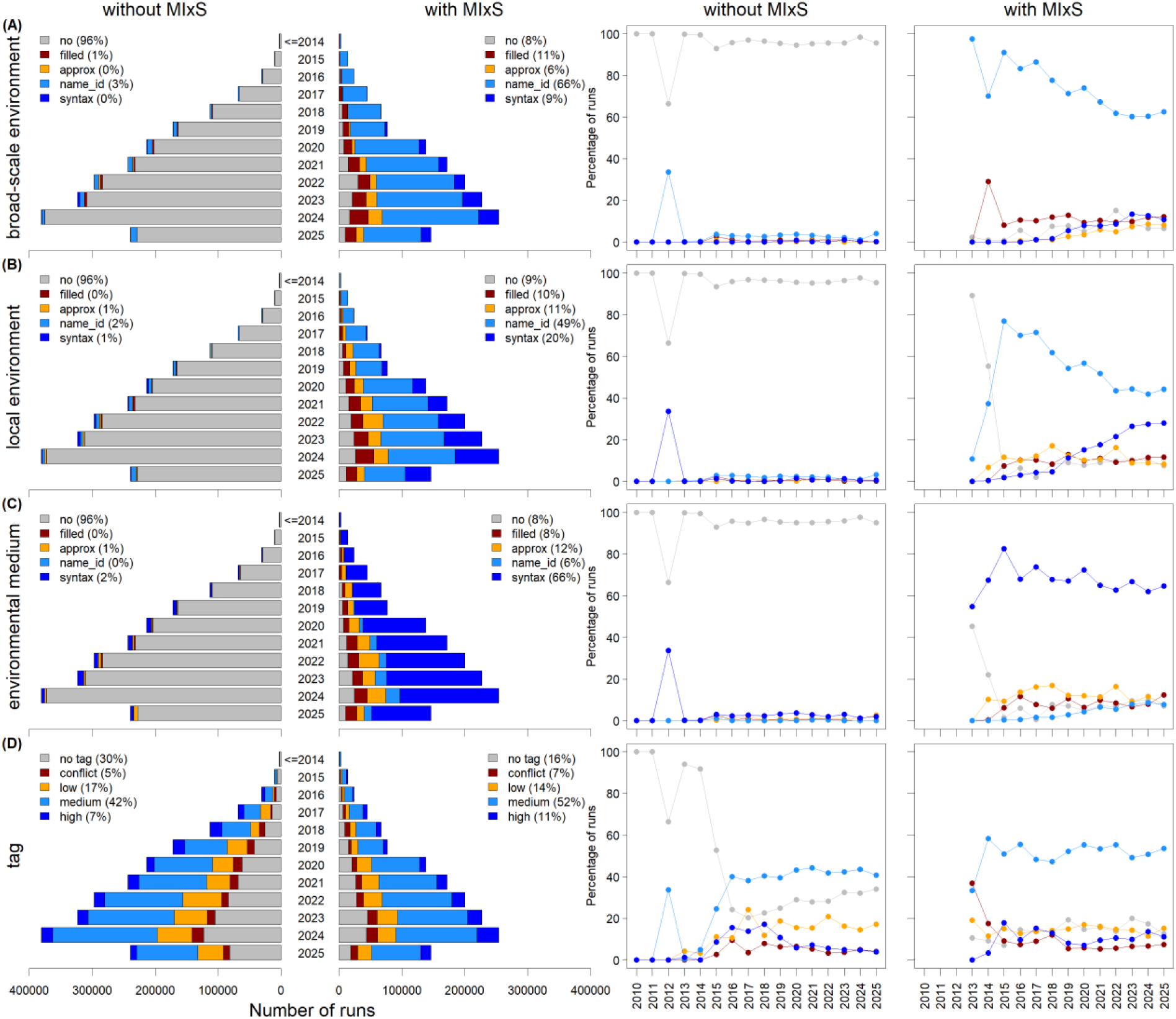
Description of the sampled environment. Values for the MIxS fields broad_scale_environmental_context (A), local_environmental_context (B), and environmental_medium (C), classified as syntax (ENVO term name and ID provided MIxS syntax-compliant), name_id (ENVO term name or ID provided verbatim), approx (matching an ENVO name after conversion to lowercase alphanumeric characters), filled (no similarity to ENVO terms), or no (no value). (D) Environment tag automatically assigned by ENA, classified as high, medium, or low confidence, or conflict (adapted from O’Cathail (2024)), or as no tag available. The bar charts show the number of runs per submission year, separately for runs submitted without and with a MIxS checklist and coloured by metadata quality category. The line charts show the percentage of runs in each category over time. Percentages in the legends refer to all years combined. Runs submitted before 2015 are pooled (≤2014) in the bar charts.

The binary indexed metadata field “environmental_sample” returned as part of the TSV search output is not element of a MIxS checklist. Values for this metadata field were not available for 99% of the runs, making it unsuitable for data mining. Instead, the automatically generated tag summarises derived information about the environmental origin of a sample. Such an assignment provided medium to high confidence prediction for more than half of the runs, classifying them as either terrestrial, coastal, marine, brackish, or freshwater (Figure 4D).

### Amplified region

Missing information about which DNA region was targeted by the amplicon sequencing approach is one of the main obstacles for data reuse. While slightly higher for submissions using a MIxS checklist, the percentage of runs with metadata provided for “target_gene” and “library_construction_protocol” in the TSV output remained low at 1.7% and 4.2%, respectively (Supplementary Table 1), and only 0.4% of all runs provided a value resembling a primer sequence in those metadata fields (Figure 5A). For the majority of the remaining runs, the non-indexed fields in the sample and experiment XML potentially contained some indication about which amplicon was sequenced, including a nucleotide sequence for 42% of those cases. However, putative information about the amplified region in the sample XML and experiment XML was provided in 241 and 24 metadata field names, respectively, that did not follow a controlled vocabulary or syntax at the level of the field name or its values. As such, the metadata is not interoperable in the machine-readable sense. A closer look at the experiment XML based on a haphazard manual check further indicated that the presented numbers likely constitute a best-case scenario, since metadata fields such as “DESIGN DESCRIPTION” did not always contain information about the amplicon. Interestingly, the use of a MIxS checklist improved the availability of TSV-, but not XML-level metadata. The fraction of runs with potential amplicon information in the XML has been increasing since 2015. Consistent with fewer MIxS-compliant submissions through NCBI, runs submitted through this entry database primarily provided putative amplicon metadata at XML level. GFBio had promoted the provision of “target_gene” at sample level until 2021, resulting in 20% of runs submitted until that time containing amplicon metadata in the TSV, although not with sequence information. In 2022, the policy changed to represent metadata about PCR conditions, which are part of the library preparation, at experiment level. Since then, no more runs were submitted with TSV-level amplicon metadata. Instead, a marked increase in the proportion of runs with amplicon metadata in the XML was observed, resulting in overall lower proportion of runs without any primer information. Compared to data sets submitted without the use of this brokerage service, the proportion of runs with a putative primer sequence in the XML was significantly increased (Supplementary Table 1) reaching almost 50% of all submitted runs.

**Figure 5:**
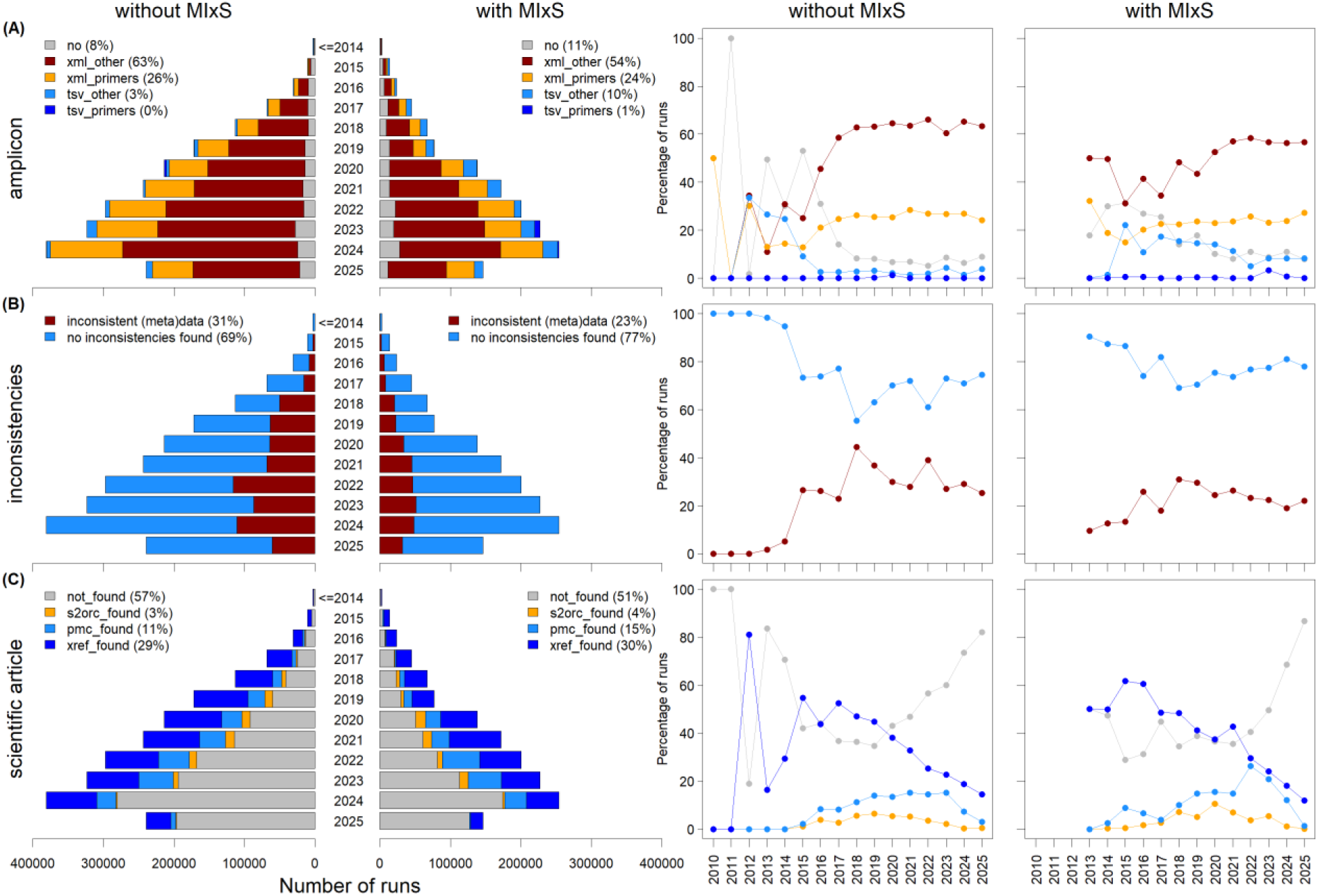
(A) Information about the amplified region in the TSV output or the sample and experiment XML, distinguishing values that contain a primer sequence (*_primers) from other values (*_other). (B) Runs with contradictory metadata. (C) Discoverability of an associated scientific article, assessed sequentially via the ENA XREF API (xref_found), a search of the PMC corpus (pmc_found), and a search of the S2ORC corpus (s2orc_found). The bar charts show the number of runs per submission year, separately for runs submitted without and with a MIxS checklist and coloured by metadata quality category. The line charts show the percentage of runs in each category over time. Percentages in the legends refer to all years combined. Runs submitted before 2015 are pooled (≤2014) in the bar charts.

### Contradictory metadata

In a large-scale assessment of metadata quality as presented in this study, it is difficult to evaluate the correctness of the provided information beyond its presence and syntax-compliance. To gain a (most likely under-)estimate of the frequency at which values in different metadata fields contradict each other, we conducted a rudimentary sanity check based on mutually exclusive scenarios during data generation: (1) DNA cannot originate from a single genome if a mixed sample was processed; (2) paired-end data require exactly two read files if it was uploaded correctly; (3) PCR is the only viable library selection method to generate amplicons from (meta-)genomic material. Even with those minimal criteria for consistency, the overall proportion of inconsistent entries was 28% (Figure 5B), primarily caused by data sets being assigned “GENOMIC” as library source despite the NCBI taxon ID labelling them as metagenomic, while violations of the other two criteria constituted the minority. MIxS-compliant submissions exhibited a significantly reduced fraction of inconsistencies (Supplementary Table 1), although still at 23% overall, with a slightly decreasing trend since 2018. Compared to NCBI, ENA (and to lesser degree DDBJ) submissions showed fewer inconsistencies (Supplementary Table 1), although runs with inconsistent metadata submitted through NCBI have also been decreasing since 2018 from 43 to 24%. While data sets brokered by GFBio were not exempt from inconsistencies, only 6% of runs were affected.

### Discoverability of associated scientific article

Metadata from the ENA portal API is convenient but often still limited, so linking a data set to its associated scientific article, where one exists, can provide valuable additional information. The ENA XREF API offers automated access to associated publications indexed on PubMed and Europe PMC. However, only 30% of runs could be linked to a scientific article via XREF (Figure 5C). Searching the PubMed and S2ORC corpora directly linked an additional 16% of runs. This number may constitute an underestimate since our approach was biased towards Open Access publications. On average, MIxS-compliant submissions had fewer runs without an associated article, although the effect was minimal and trends over time were comparable to runs from submissions without MIxS. Specifically, the fraction of runs with scientific articles linked via XREF has been decreasing since 2015. Approximately until 2022, this development was to some degree buffered by an increasing fraction of runs linked to a scientific article via the PubMed and S2ORC corpora, after which the fraction of runs without an associated scientific article increases rapidly. These trends may arise from multiple reasons: there may be technical issues with XREF regarding the accessibility of the ENA study accession in the article, a change in the fraction of full articles available through PubMed and EuropePMC^39^, an offset in the publication time between the data and the article, or a stronger mobilisation of data for which no publication in form of a scientific article is intended.

Taken together, the number of runs with standardised machine-readable metadata for all investigated metadata fields, which in themselves only constitute a fraction of recommended checklist fields, is shockingly low: Only 2% of MIxS-compliant and 0.1% of non-MIxS runs met our criteria for machine-readability, with an overall fraction of 0.7% (Figure 6). The largest fraction of runs contained human-readable metadata, i.e. non-standardized values in the investigated metadata fields or an associated scientific article. This fraction has been decreasing since 2019 for non-MIxS and since 2021 for MIxS-compliant data sets, primarily linked to the availability of an associated article. Over all years and across the whole metadata set, these trends resulted in a total fraction of 27% of runs with missing or inconsistent metadata (or no fastq files) and no retrievable additional information to remedy that.

**Figure 6:**
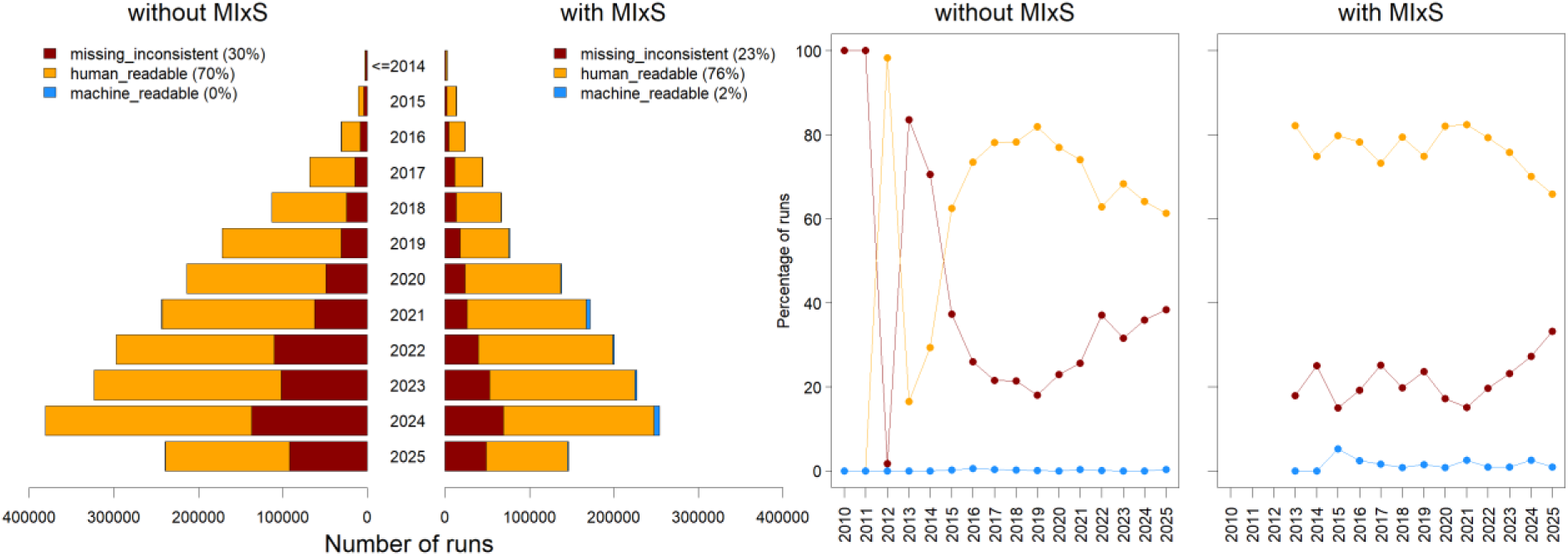
Overall metadata quality per run, combining all assessed fields: machine-readable (syntax-compliant and consistent metadata in the TSV output), human-readable (metadata available in TSV or XML format and/or a linked scientific article), and missing/inconsistent (missing or contradictory metadata, or no fastq files, without a linked article). The bar charts show the number of runs per submission year, separately for runs submitted without and with a MIxS checklist and coloured by metadata quality category. The line charts show the percentage of runs in each category over time. Percentages in the legends refer to all years combined. Runs submitted before 2015 are pooled (≤2014) in the bar charts.

### Standard compliance of sequence read data

Of the 39 studies inspected in Molari et al.^31^ that sequenced the V3-V4 hypervariable region of the bacterial 16S rRNA gene, only eight were submitted as required (Figure 7A). The majority (28 studies) did not remove the primer sequences, eight studies contained already merged sequences, and one study even provided the sequencer output prior to demultiplexing (sample barcode information had to be obtained from the author). In the second example, Riedinger et al.^30^ effectively reused paired-end Illumina amplicon sequencing data from 710 studies, of which we assessed 343 that sequenced the V3-V4 hypervariable region of the 16S rRNA gene. Only 26 studies (8%) provided their data in compliance with submission requirements unmerged and without primers, while the large majority (300 studies; 87%) still contained primers (Figure 7B). The remaining 17 studies, contained merged reads provided in a single fastq file, either primer-clipped (11 studies) or with primers included in the sequences (6 studies).

**Figure 7:**
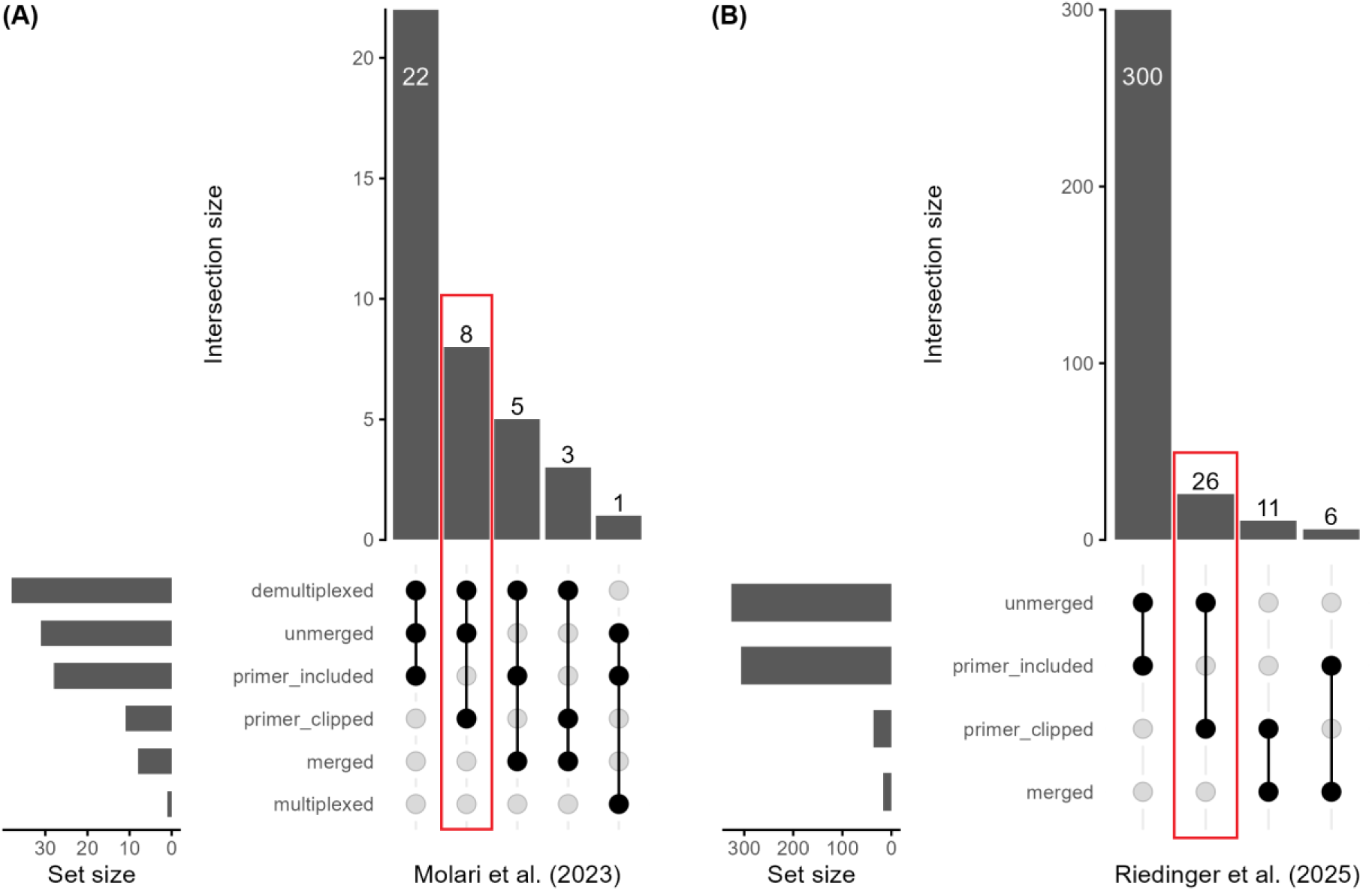
Upset plot showing the number of ENA studies reused in two case studies (A: Molari et al.^31^; B: Riedinger et al.^30^) containing fastq files submitted at various levels of pre-processing. Submission guidelines require demultiplexed, primer-clipped, unmerged sequencing read data (red square).

These use cases showed that even if metadata that enables the findability and accessibility of the data is provided, the sequence read data itself may often not be submitted as required, making manual checks mandatory and limiting the reusability of the data.

## Discussion

All investigated data sets met the criteria for findability and accessibility since those were a prerequisite of conducting this study. Data sets not discoverable by our query were by necessity excluded from the assessment. Furthermore, while accessibility of the metadata did not guarantee the accessibility of the data itself, as part of the runs contained no sequence files (0.25%), these data sets can still be classified as accessible in accordance with Wilkinson et al.^6^. However, interoperability, and therefore reusability, remained a challenge. Overall, our results revealed a high variability in data and metadata interoperability and reusability for metabarcoding read data, depending on the investigated metadata field. In combination, our results were sobering: Readily machine-readable, syntax-compliant, and consistent metadata was available for less than 1% of the investigated data sets and almost 30% exhibited deficiencies in their metadata unlikely to be resolvable to facilitate data reuse or lacked data altogether. The number of data sets being submitted each year is expected to continue to rise considering decreasing sequencing costs and the undiminished popularity of amplicon sequencing for multiple applications in molecular and microbial ecology and biodiversity research (e.g. environmental DNA studies). It is necessary to overcome the obstacles in FAIR data management that have resulted in the current state of metadata quality. Data reusability is required for the sustainable research practices allowing data to accrue added value by becoming a common good. The importance of taking action is thereby not restricted to basic research. Metabarcoding data plays an essential role in biodiversity policy instruments, such as the Global Biodiversity Framework^3^, where improving data availability is highlighted as one of its main targets for 2030 (target 21).

### Interoperability: syntax and semantics

The metadata fields that are mandatory for all submissions, i.e. “collection_date” and “country”, were syntax-compliant for the large majority of runs. However, despite superficial syntax compliance, an alarmingly large fraction of values for “collection_date” exhibited a low interoperability by not including day resolution. The flexible syntax implementation, necessary for data sets where uncertainty regarding the collection date does exist, was likely exploited by data submitters providing imprecise metadata (year or month only) although a precise date or date range would have been possible as confirmed by manual cross-checks with the associated publication. The controlled vocabulary used for “country” is based on the <u>standard country or area</u> <u>codes for statistical use</u> of the UN and the <u>CIA World Factbook</u>. Deviations from this vocabulary or allowed values for missing data were rare and mainly restricted to no “country” information being provided. As such, investigated data sets are likely highly interoperable with other data resources that are also referring to UN standards, although interoperability could be improved by assigning PIDs to the terms in the controlled vocabulary^40^. Correct identification of the country of origin for a sample from which sequence information was derived and its collection date are essential for equitable <u>Access and Benefit Sharing</u> and therefore highly relevant beyond a purely scientific context.

For a data set to be reusable, it has to contain sufficient metadata to place it within the context of other data sources. “Country” information is insufficient for this purpose. For instance, latitude, longitude, vertical and time coordinates are required to link sequence-based biodiversity observations to the Ocean Biodiversity Information System (<u>OBIS</u>), the Global Biodiversity Information Facility (<u>GBIF</u>), and other platforms for biodiversity assessment^41–43^. Furthermore, they are needed to retrieve additional data about the sampled environment from other data repositories, e.g. Pangaea^44^, Copernicus^45^ (remote sensing), or to interpolate sequence-derived biodiversity data with gridded data, such as derived from Earth System models^46^. In some cases, concerns about disclosing personal data and the location of endangered species^47^, or the suitability of coordinates for laboratory facilities and control samples in an experimental setting, constitute valid reasons to not provide exact coordinates. However, metadata still needs to be provided and the respective metadata fields should not be left blank, since all these cases are represented in the controlled vocabulary for missing values^27^.

While latitude and longitude were readily interoperable (if provided), the situation for vertical coordinates was more complicated with regard to both the definition of the metadata field and its syntax. For instance, “depth” is semantically confounded, since it is defined relative to the closest surface of a sample instead of a fixed reference like geodetic surface. As such, the depth of a sample will mean something different depending on whether it was collected in soil, water, or sediment. Furthermore, the documentation about the syntax for depth values is contradictory between the metadata standard and its implementation in submission checklists. According to MIxS, the unit is part of a <u>depth</u> value with meters being recommended but not enforced^48^. In the ENA submission checklists, the syntax is ambiguous regarding the inclusion of the unit, with both m or mm recommended, leaving the choice up to the user^17^. According to the ENA portal API documentation, depth is only defined in meters for water, but not for sediment and soil samples, which lack such a definition^18^. The brokerage service GFBio requires no unit to be part of the value, but expects values in meters^49^, which are then included as value/unit pair in the sample XML. However, depth values are not represented consistently this way in the sample XML, with the unit frequently part of the “value” without there being a separate “unit” attribute and no restrictions as to which units are allowed, including non-SI units. Consequently, the current practices for providing the vertical location of a sample do not reflect actual usage requirements. Such information is among the most basic metadata, but already problematic both in terms of definition and use.

Coordinates in space and time may not always be available or reliable, and even if they are, they may not unambiguously allow to infer the sampled environment. Correct identification, e.g. of samples from shorelines, host organisms, laboratory or experimental settings, or other environments, where location may be confounded by additional factors, often requires further information that is difficult to extract from available metadata. The issue starts with the definition of “environmental”: Commonly, in ecology the term is used to describe samples originating from unmanipulated environments, e.g. field but not laboratory-sourced experimental samples. However, within the context of sequence archives, the term (and metadata field) “environmental_sample” refers to whether the nucleic acid was extracted from a bulk (mixed) sample as opposed to a single organism or strain^50^. Therefore, mixed samples from a laboratory experiment would also be described as “environmental_samples” under the INSDC definition. This apparent confusion about the use of this metadata field makes it unsuitable to guide data reuse. When derived from latitude and longitude alone, the metadata field “tag” is strongly affected by the above-mentioned bias regarding the inference of the sampled environment from the location. This bias is alleviated by using another metadata field that is mandatory: the NCBI taxon ID. While this may be functional, it also represents an improvised solution. That the taxon ID includes semantics about the environment is incidental, and it is ironic that a metadata field that was never intended for this use is now among the most meaningful descriptors for environment - even more so since the second most frequent taxon ID is uninformative (metagenome).

MIxS introduced a trilogy of metadata fields to describe the environmental context of a sample using the environmental ontology (ENVO^10,11^): broad and local scale environment, and environmental medium. However, this has not resulted in a sufficient level of interoperability and reusability of sequencing data sets. There are discrepancies between the MIxS schema and its implementation in the INSDC submission checklists, as well as between the schema and observed common practice. Even in MIxS compliant submissions, the metadata fields describing the environment were often not used as intended. In part, this situation may have arisen from inconsistencies in the documentation about metadata requirements provided by separate resources: the ENA documentation specifies free text^17^, whereas MIxS specifies the use of a controlled vocabulary and syntax with cardinality depending on the metadata field^48^. This lack of congruency is caused by different institutions responsible for defining the MIxS metadata standard (GSC) and implementing it (INSDC). Consequently, not all GSC recommendations are adopted into submission checklists. In such cases, it is the responsibility of the user (or broker) to adhere to the more stringent standard in compliance with its original description. A potential reason for the reluctance of INSDC to enforce MIxS syntax may lie in its lower user-friendliness, resulting in a trade-off between interoperability and incentives for users to provide any information at all, and in the additional strain this would put on their helpdesks. Hopefully, recommender tools (BiodivPortal^51^) and advances in LLMs^52,53^ can overcome this trade-off in the future. Furthermore, validation scripts, such as DataHarmonizer^54^, may ensure metadata FAIRness when run prior to data submission to an INSDC database.

### Misconceptions by data submitters and consumers

Despite available standards and checklists, if and how metadata are provided (and how metadata are utilized for data mining) is to a large degree subject to interpretation by the respective data submitter and data consumer. Available documentation may be ignored if the meaning of a metadata field seems apparent, or misunderstood if it is incomplete or too technically phrased. Metadata fields listed as recommended may not be questioned, whether they are actually required for data reuse depending on the type of sequencing approach that was used to generate the data.

Early versions of MIxS implemented so-called conditionally mandatory metadata fields, which by definition were mandatory if a condition was met, resulting, e.g., in “target_gene”, “target_subfragment”, “pcr_primers” as mandatory for all metabarcoding data sets. This distinction is not represented in newer MIxS versions anymore, although the need for this information has not changed^55^. The implementation of these three MIxS metadata fields at INSDC results in further problems that contributed to the observed low interoperability, if values were provided at all. Among the three, only “target_gene” is indexed as a searchable and returnable sample-level metadata field. However, information about the amplification step during library preparation is a feature of the sequencing experiment, and not the sample. This distinction is crucial because more than one experiment may be associated with the same sample. However, since MIxS is implemented at sample level, submitters are being misled and metadata may be incorrectly displayed despite best intentions. For instance, an incorrect value for “target_gene” may be inherited by the ENA portal API for raw read data of samples used for metabarcoding with multiple marker genes. Conversely, data creators may submit experiments of the same sample under different sample accessions to ensure that PCR information is displayed correctly.

The current ENA experiment XML schema includes two metadata fields that may hold PCR information: “LIBRARY_CONSTRUCTION_PROTOCOL” and “TARGETED_LOCI”. Neither field is mandatory, and neither enforces a controlled vocabulary or syntax. Of the two, “LIBRARY_CONSTRUCTION_PROTOCOL” seems less suitable than “TARGETED_LOCI”, with the former intended for a general description of the library preparation and the latter requesting specific information about the “gene targeted by the sequence”^56^. “TARGETED_LOCI” has been recently implemented at GFBio for brokering MIxS-compliant metadata on “target_gene”, “target_subfragment”, and “pcr_primers” at experiment level. However, it is not yet possible to retrieve this information parsed as TSV via the ENA portal API. Despite recent endeavours to extract this information from the sequencing data itself^57^, the correct identification of the amplified region without any respective metadata remains error-prone, cumbersome and computationally expensive. Complete and correct metadata entries, using a standardized format or even controlled vocabulary preferably in accordance with existing ontologies, and easy access to that information in human- and machine-readable format would drastically reduce computational and man-power requirements for post-deposition data curation and reuse.

We further noticed inconsistencies in the provided metadata that may have resulted from incorrectly interpreted submission instructions or oversights during the data submission process. Certain metadata values are mutually exclusive. For instance, the same data set cannot be described with both “library_source” METAGENOMIC and “library_selection” RT-PCR, since the latter implies that RNA was used to construct the sequencing library. Such contradictions are often unresolvable for the data consumer without additional information that may not be readily available or machine-readable, hampering or excluding data reuse. Differentiated data submission pipelines that restrict permitted values depending on a mutually exclusive meaning or extended plausibility checks during submission validation may help to prevent such errors.

Another example of contradictory metadata also concerns the field “library_source”. We noticed that sequencing data sets whose NCBI taxon ID clearly identified them as originating from metagenomic material, were declared as “library_source” GENOMIC. Since metabarcoding and amplicon sequencing are not synonyms for metagenomics as a methodological approach, it may be counterintuitive for a submitter to include the value METAGENOMIC in their metadata. However, the meaning of the metadata field “library_source” is not related to the sequencing approach, but the DNA extract that was processed for sequencing. Therefore, the template for the PCR during library construction was metagenomic and should be declared as such. Further confusion may have arisen when analysing host-associated microbiomes, assuming that the genomic material of the host organism may have been confused with the “library_source” of the metabarcoding experiment, which should refer to the microbiome, which by definition constitutes metagenomic material. A data consumer is unlikely to consider data sets labelled as “library_source” GENOMIC if they are interested in metabarcoding data, resulting in underestimation of available data sets. In an even worse scenario, a submitter may have used the taxon ID of the host to describe data derived from its microbiome. As noted previously, the taxon ID is used to infer the environmental origin of a sample and among the most informative metadata fields to select data for reuse. A false claim about the taxonomic affiliation of the sequences in a data set is very difficult to identify without a full re-analysis of the sequence reads, which is why we could not quantify its occurrence here, and may result in severe misinterpretation and incorrect data reuse in downstream analyses. Instead, a diligent submitter should use the MIxS metadata field “host_taxid”, while a member of <u>organismal metagenomes</u> should be selected for NCBI taxon ID of the sample.

Independent of our analysis of available metadata, the submitted data (fastq files) did not always meet the required criteria according to the submission guidelines. The presence of more or less than two files available for download (“fastq_ftp”) for paired-end read data indicated errors in read pair matching at ENA that may have resulted from corrupted fastq files or by wrongly paired files (although the data itself may have been good). Such errors are unrecoverable without substantial manual curation and the data consumer has to carefully decide whether to exclude potentially good data or risk including corrupted data. Furthermore, sequences generated as paired-end may have been submitted with metadata declaring “library_layout” single if reads were merged prior to submission despite instructions to the contrary. Our use case examples revealed that such cases indeed occurred quite frequently. We further showed that the majority of read data did not comply with the requirement that artificial reads (including primers) must be removed prior to submission^19^. Additional to deficient metadata, we therefore suspect that a large proportion of raw read metabarcoding data not been archived correctly according to the INSDC submission guidelines. In the case of primer presence, the proportion of read files still containing primers was so high that it may be more feasible to adjust the submission requirements than to convince the submitter community to change their practices.

Within the scope of this study, our analysis was limited to metadata and data inconsistencies that were fairly easy to assess. Other issues, which we did not explore in more detail here include (1) duplicated data submissions: we observed identical submitted md5sums for 24,968 runs (out of the 422,066 runs submitted via ENA), which seems highly unlikely to occur randomly; (2) undifferentiated metadata entries, presumably resulting from copy paste operations in free text descriptors: e.g. control samples being described as environmental, experiments described as both 16S and 18S, uninformative and uniform descriptions for all samples in a study; (3) excel auto-fill errors that convert stationary coordinates into a transect or increment names ending with a number; (4) wrong decimal separator for numeric environmental variables, e.g. temperature, salinity; (5) errors in the SRA accession number referenced in the scientific article. Such inconsistencies are especially difficult to track and often only detectable after a manual check of the data, metadata, and associated publication. Still, superficial, haphazard checks revealed a worrisome situation that emphasizes the need for more stringent validation routines prior to data submission. As it stands now, more than a quarter of archived metabarcoding data is unsuitable for reuse and therefore likely lost to future generations.

### The way forward

Our study adds another facet to the increasing body of work illustrating the deficits in nucleotide sequence databases^21–23^ with drastic consequences: Terabytes to Petabytes of data may not be readily interoperable and reusable, severely limiting their added value, long-term impact, and future relevance. While the number of tools, platforms, and tutorials to inform and facilitate sustainable data management has increased rapidly in the recent past^58–61^, data literacy remains an issue and trends in our analysis do not indicate sufficient improvement. This shows that placing the task of improving FAIRness solely on data submitters will not yield a satisfactory outcome. Primarily a shift in the recognition and scientific value system is required to provide incentives for proper data archival and publication, including standardized metadata, to enable long-term reuse of the data^60–62^.

The MIxS metadata standard has been established more than a decade ago^9^. Since then, there have been multiple calls to use MIxS^63–65^ and MIxS has been integrated into dedicated custom checklists (often with stricter implementation), e.g. FAIReDNA^55^ as a response to the widely recognised need of the eDNA data ecosystem for improvements centred around FAIR. A recent roadmap provides guidance for how to achieve this specifically for aquatic eDNA^66^. Despite the issues with the MIxS implementation at INSDC databases outlined previously, our analysis showed that the use of a MIxS checklist significantly increased syntax compliance and therefore interoperability. This effect was closely linked to the INSDC database through which the data were submitted since fewer submissions used MIxS at NCBI. While only a small percentage of runs were submitted via GFBio, the use of this brokerage service had a profound positive effect on interoperability, fundamentally improving metadata quality and therefore data reusability.

Brokerage services are essential to ensure high (meta)data FAIRness and act as intermediary between the data submitter and the database. As an additional checkpoint, they constitute a fulcrum where automatic validation tools and plausibility checks may be most effective, contributing expertise in data management and bioinformatics, and hosting the computational resources required for implementing such tools. Thereby they reduce both the load placed on each data submitter as well as the database helpdesks. With an increasing and diversifying amount of data, metadata requirements to achieve FAIRness are going to become more complicated. Data brokers and curators are desperately needed and adequate funding is required to ensure sustainable service provision without them being overwhelmed (supported by UN Ocean decade^67^, worldFAIR^68^, CODATA^69^).

Recent advances in AI technologies open another avenue for improving metadata FAIRness, both for future data submission as well as for mobilising existing data. The verdict presented here is largely based on metadata fields that require controlled vocabulary according to a clearly defined syntax. AI tools make free text metadata fields and natural text, e.g. the scientific article describing a data set, accessible for extracting structured information^52^. Such approaches hold enormous potential to reduce the manual time investment otherwise necessary for data curation, thereby increasing the data volume available for reuse. Indeed, Ikeda et al.^52^ recently demonstrated opportunities and limitations of using AI technologies to improve BioSample fields’ values. However, in general the application of AI technologies still requires a data set to be identified first as potentially suitable for reuse in a specific context, which may not always be the case as outlined previously, and a publication linked to the data set if available. Our results show that the latter is not reliably implemented yet. For instance, the ENA XREF API is not able to retrieve all possible links between data sets and publications present in PMC/PubMed. The metadata exchange landscape still needs to mature in its intersection with data on the web as bibliographic (meta)data typically does not address machine-readable data availability statements^70^.

## Conclusions

Many of the necessary developments outlined above require time. In the meantime, the volume of metabarcoding sequencing data keeps growing without improvements in FAIRness. It is crucial to avoid further delays in improving (meta)data quality by making better use of existing standards and current tools and services for sequence data management. In the following we provide a (non-exhaustive) list of suggestions to address some of the most acute deficits of data and metadata FAIRness for metabarcoding sequencing data. They were selected because we think that they are comparatively easy to implement and will have a potentially large positive impact on the research fields that rely on such data, as well as on derived applications in biodiversity, conservation policy and management strategies.

### Suggestions @researchers

- Make use of existing checklists and data brokerage services, and use checklists beyond mandatory parameters. For instance, the MIxS parameters “target_gene”, “target_subfragment”, and “pcr_primers” should be supplied for all sequencing read data that was generated with library_selection=“PCR” AND library_strategy=“AMPLICON”.
- Enter data diligently and according to the specified format to facilitate interoperability. It is not only important **that** (meta)data is archived, but also **how**.
- Whenever possible, use ontologies and actively contribute to the improvement of ontologies by suggesting so far missing terms to the ontology developers. Many ontologies, like ENVO, have a low-threshold for suggesting a new term, e.g. opening an issue on GitHub, and provide guidance on using ontology terms in the context of MIxS^71^.
- Be specific in free text descriptions. Differentiate between samples in their description, provide environmental or experimental conditions. Such free text information is accessible to AI tools.
- Update data submissions if additional information (manuscript DOI, accession numbers of related data sets) becomes available or errors in the metadata are noticed.
- Account for the costs of FAIR data management in project proposals by requesting additional funding for data technicians and data managers for the successful implementation of data management plans.

### Suggestions @research institutions

- Invest in capacity development and training of early career researchers to avoid incorrect data (and associated metadata) submissions due to inexperience, and raise awareness for the persistently high value of FAIR data. Data archiving is not a trivial task, and it is at least as important as manuscript publications.
- Incentivation of good data management through recognition of data publications towards career progression metrics.

### Suggestions @reviewers

- Review the submitted data and metadata for a scientific manuscript as thoroughly as the manuscript text. FAIR data archiving should be as important a criterion for manuscript publication as scientific soundness. Peer-review can be used to distribute the burden of data curation. Mechanisms as suggested in Hug et al.^72^ can be used to alleviate the fear of scooping for data sets made public prior to manuscript acceptance to grant reviewers access when no dedicated reviewer access is available.

### Suggestions @publishers

- Make reviewing the data and metadata a mandatory aspect of peer-review that is given equal importance in editorial decisions as the scientific soundness and reproducibility of the methods section of a manuscript.

### Suggestions @databases & brokers

- Implement automated plausibility checks for data consistency, specifically related to the use of controlled vocabulary (ontologies) and mutually exclusive metadata values or make such tools easily available for pre-submission checks. Coordinate their development and implementation with data brokers.
- Harmonize the documentation about the parameters that use a controlled vocabulary (ontologies) across the different resources (MIxS, ENA) by choosing the more stringent standard.
- Expand the list of indexed metadata (MIxS) parameters in a concerted effort with the scientific community to promote stronger adherence to more extensive checklists and standards.
- Provide experiment metadata checklists that include relevant MIxS fields. MIxS is not only intended for sample checklists, but covers sample and experiment metadata. In this context we note the cross repository working group at the Global Alliance for Genomics and Health (<u>GA4GH</u>) and encourage experimental field alignment with MIxS.

## Data Availability

No new data, i.e. no non-code output, was generated for this study. The metadata quality assessment tables, computed with the code stated below and used to generate the presented figures, are available in Supplementary Table 2.

## Code Availability

The scripts used in the metadata analysis are available on https://github.com/chassenr/ENA-metadata. The README outlines the analysis workflow, the directory structure created during the analysis, and which scripts to use at which analysis step.

## Acknowledgments

Many thanks to Anna-Adriana Anschütz for her persistence in helping me solve R plotting issues.

## Author Contributions

Conceptualization: C.H., P.L.B., M.M., V.H., I.K.

Methodology: C.H., J.M., R.M.

Formal analysis: C.H., J.M., T.P., D.R.

Visualization: C.H., J.M.

Writing - Original Draft: C.H., J.M.

Writing - Review & Editing: All authors

## Supplementary information

**Supplementary Figure 1:**
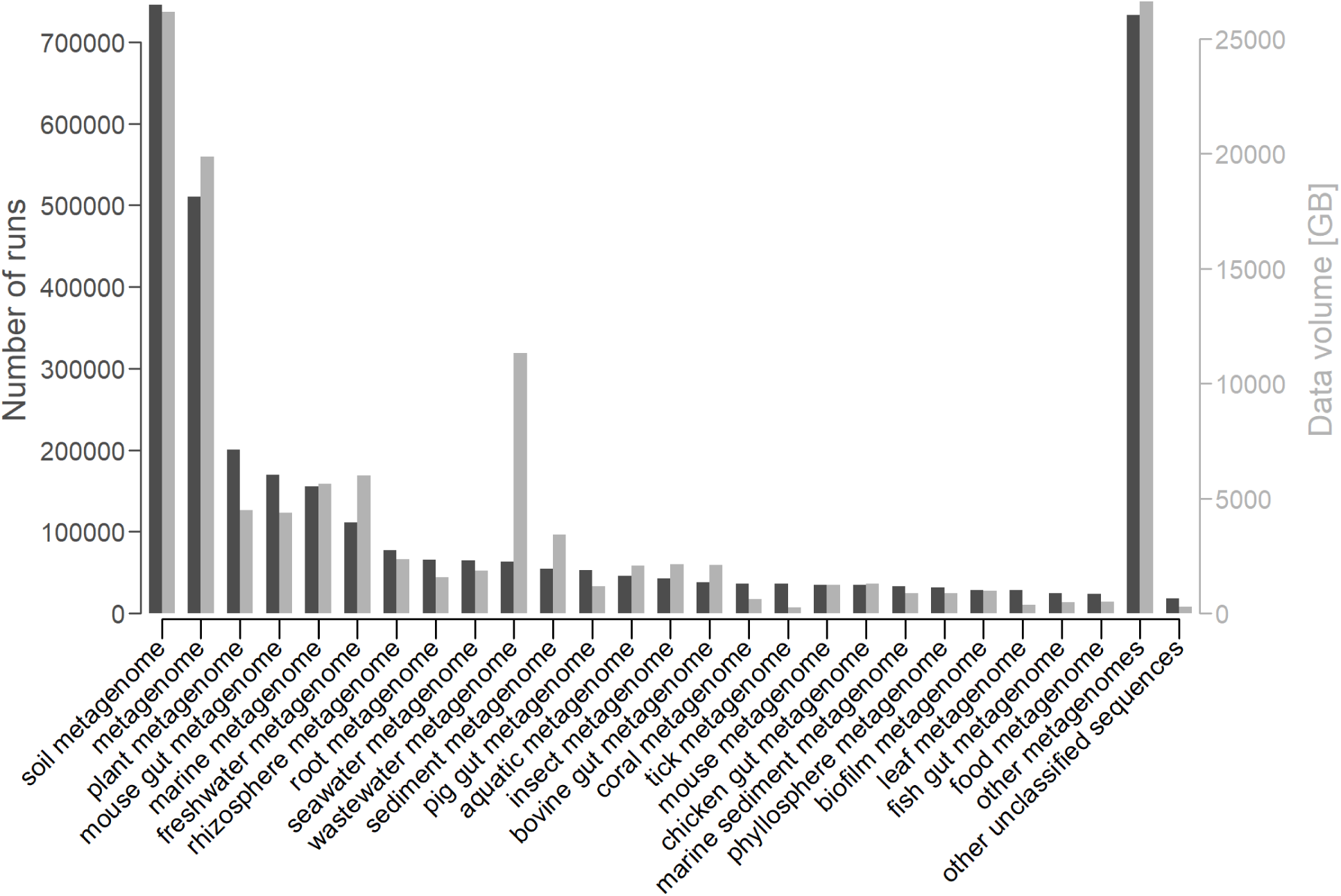
Taxonomic composition of the analysed data selection, sorted decreasingly by number of runs (left axis, dark gray). After the top 25 most frequent NCBI taxon IDs, the remaining taxa were grouped into “other metagenomes” (children of taxon ID 408169). “Other unclassified sequences” then captured all NCBI taxon IDs not represented in the previous categories. Corresponding data volume is shown on the right axis (light gray).

